# The SARS-CoV-2 Envelope and Membrane proteins modulate maturation and retention of the Spike protein, allowing optimal formation of VLPs in presence of Nucleoprotein

**DOI:** 10.1101/2020.08.24.260901

**Authors:** Bertrand Boson, Vincent Legros, Bingjie Zhou, Cyrille Mathieu, François-Loïc Cosset, Dimitri Lavillette, Solène Denolly

**Affiliations:** CIRI – Centre International de Recherche en Infectiologie, Univ Lyon, Université Claude Bernard Lyon 1, Inserm, U1111, CNRS, UMR5308, ENS Lyon, 46 allée d’Italie, F-69007, Lyon, France; Université de Lyon, VetAgro Sup, Marcy-l’Étoile, France; Institut Pasteur of Shanghai, Chinese Academy of Sciences; University of Chinese Academy of Sciences; CAS Key Laboratory of Molecular Virology & Immunology, Institut Pasteur of Shanghai Chinese Academy of Sciences Pasteurien College, Suzhou University, Jiangsu, China

## Abstract

The severe acute respiratory syndrome coronavirus 2 (SARS-CoV-2), a β-coronavirus, is the causative agent of the COVID-19 pandemic. Like for other coronaviruses, its particles are composed of four structural proteins, namely Spike S, Envelope E, Membrane M and Nucleoprotein N proteins. The involvement of each of these proteins and their interplays during the assembly process of this new virus are poorly-defined and are likely β-coronavirus-type different. Therefore, we sought to investigate how SARS-CoV-2 behaves for its assembly by expression assays of S, in combination with E, M and/or N. By combining biochemical and imaging assays, we showed that E and M regulate intracellular trafficking of S and hence its furin-mediated processing. Indeed, our imaging data revealed that S remains at ERGIC or Golgi compartments upon expression of E or M, like for SARS-CoV-2 infected cells. By studying a mutant of S, we showed that its cytoplasmic tail, and more specifically, its C-terminal retrieval motif, is required for the M-mediated retention in the ERGIC, whereas E induces S retention by modulating the cell secretory pathway. We also highlighted that E and M induce a specific maturation of S N-glycosylation, which is observed on particles and lysates from infected cells independently of its mechanisms of intracellular retention. Finally, we showed that both M, E and N are required for optimal production of virus-like-proteins. Altogether, our results indicated that E and M proteins influence the properties of S proteins to promote assembly of viral particles. Our results therefore highlight both similarities and dissimilarities in these events, as compared to other β-coronaviruses.

**Author Summary:** The severe acute respiratory syndrome coronavirus 2 (SARS-CoV-2) is the causative agent of the COVID-19 pandemic. Its viral particles are composed of four structural proteins, namely Spike S, Envelope E, Membrane M and Nucleoprotein N proteins, though their involvement in the virion assembly remain unknown for this particular coronavirus. Here we showed that presence of E and M influence the localization and maturation of S protein, in term of cleavage and N-glycosylation maturation. Indeed, E protein is able to slow down the cell secretory pathway whereas M-induced retention of S requires the retrieval motif in S C-terminus. We also highlighted that E and M might regulate the N glycosylation maturation of S independently of its intracellular retention mechanism. Finally, we showed that the four structural proteins are required for optimal formation of virus-like particles, highlighting the involvement of N, E and M in assembly of infectious particles. Altogether, our results highlight both similarities and dissimilarities in these events, as compared to other β-coronaviruses.

## Introduction

At the end of 2019, SARS-Cov-2 emerged in China through zoonotic transmission and led to the COVID-19 pandemic, cumulating to date to over 16 million cases and more that 600,000 deaths worldwide [1]. SARS-CoV-2 belongs to the β-coronavirus genus of the *Coronaviridae* family that includes SARS-CoV-1 and Middle East Respiratory Virus (MERS-CoV), which are also responsible for severe lower respiratory infections.

The main structural components of coronaviruses are the S (Spike) glycoprotein, the M (Membrane) and E (Envelope) transmembrane proteins, and the N nucleoprotein, which forms a viral ribonucleoprotein (vRNPs) complex with the 30kb-long viral genomic RNA (vRNA). The S glycoprotein is the major determinant of viral entry in target cells. The M glycoprotein is key for assembly of viral particles by interacting with all other structural proteins [2, 3], whereas the E protein is a multifunctional protein, supposed to act on viral assembly, release of virions and pathogenesis (reviewed in [4]). Specifically, through its oligomerization, E forms an ion-channel termed ‘viroporin’ [5, 6]. Even though M coordinates virion assembly, an interaction between M and E is required for the formation of viral particles [7–9].

Coronaviruses assembly and budding occur in the lumen of the endoplasmic reticulum (ER)-Golgi intermediate compartment (ERGIC) [10, 11]. To ensure their accumulation in the ERGIC, M, E and S proteins contain intracellular trafficking signals that have been identified for some coronavirus species. For example, a dibasic retrieval signal, KxHxx, found at the C-terminus of the cytoplasmic tail of SARS-CoV-1 Spike allows its recycling *via* binding to COPI [12]. Such a recycling of S may increase its chance to interact with M, which resides at the ERGIC, hence inducing S accumulation at the virion budding site.

Here, we aimed at better characterizing the interplay between S and the other structural proteins E, M and N proteins of SARS-CoV-2. Owing to its homology with β-coronaviruses, we hypothesized that some assembly mechanisms might be conserved between SARS-CoV-2 and other β-coronaviruses. Specifically, we aimed at determining how M, E and N might regulate S intracellular trafficking and maturation properties. Specifically, we also investigated the furin-mediated cleavage of S, which is not present in SARS-CoV-1 [13]. Furthermore, since SARS-CoV-1 has been proposed to induce the release of S-containing virus-like particles (VLPs), though through poorly characterized processes, we also aimed at clarifying the minimal requirement for production of S-containing VLPs for SARS-CoV-2.

## Results

### Processing of SARS-CoV-2 Spike protein is influenced by other viral proteins

We compared the expression of the S glycoprotein in VeroE6 cells upon infection with full-length SARS-CoV-2 *vs*. transfection of a S-expressing plasmid at 48h post-transfection or infection (Figure 1A). We detected both a predominant non-cleaved S form, denoted S0 (of *ca*. 180kDa,) and a cleaved form of S, denoted S2 (of around 110kDa), which is likely induced from S0 processing by furin [13], an ubiquitous protein convertase localized inside the cell secretory pathway [14]. We found that S2 and S0 appeared as doublets in transfected cells, with less intense bands appearing above the bands that co-migrated with those detected in infected cells. Interestingly, we detected *ca*. 10% of S2 in infected VeroE6 cells vs. up to 40% in S-transfected VeroE6 cells (Figure 1A-B), suggesting that some other viral proteins may influence S cleavage rates. In addition, we observed in SARS-CoV-2 infected cells a second form of S2, denoted as S2*, migrating faster than S2 (around 90-100kDa), which was not detected in S-transfected cells (arrow in Figure 1A), hence suggesting also that other viral proteins influence the maturation of S2.

**Figure 1.**
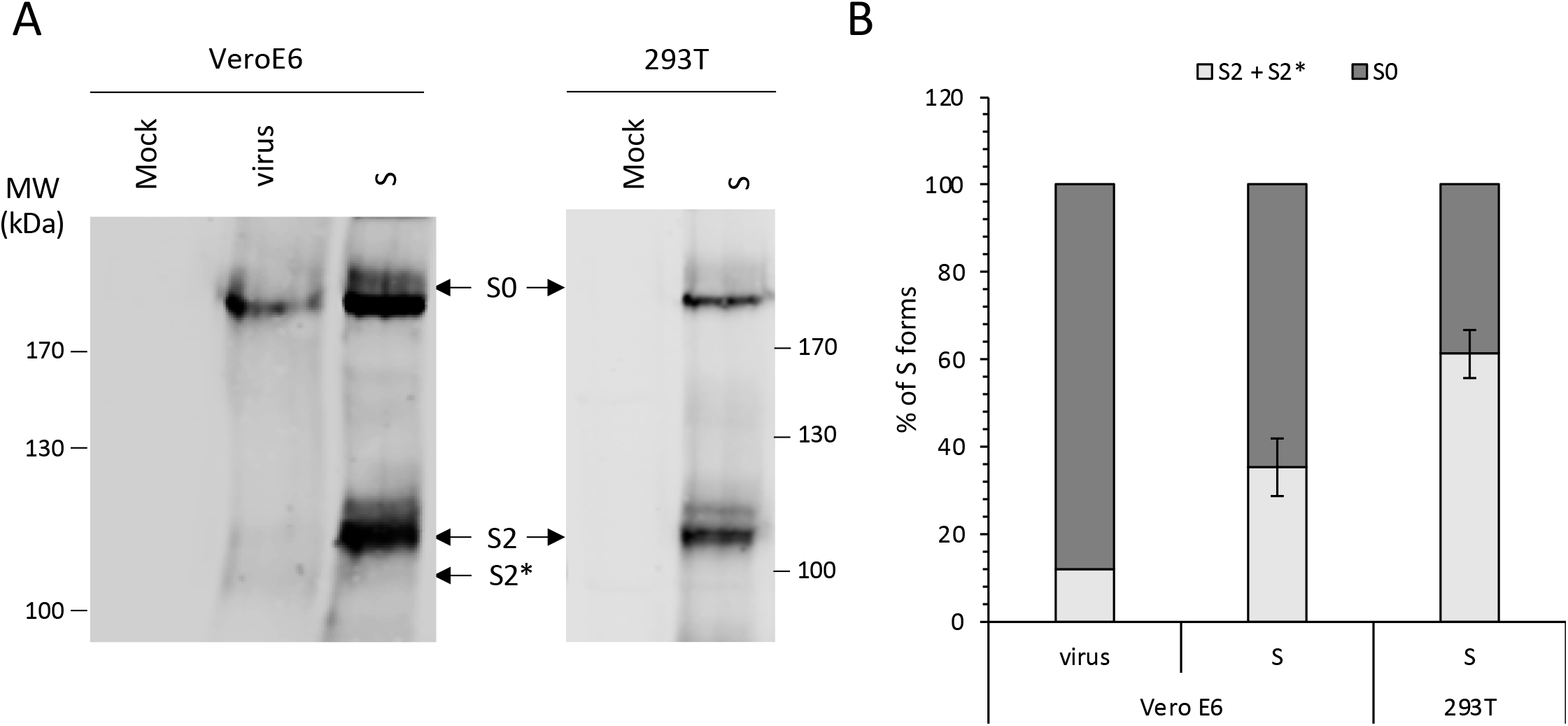
Processing of SARS-CoV-2 Spike protein is influenced by other viral proteins. **(A)** Representative Western blot of cell lysates from VeroE6 infected cells (virus) or transfected with plasmid encoding S (S) and 293T transfected with the same plasmid at 48h post infection or transfection. The arrows represent S0, S2 and S2* forms. **(B)** Quantification of proportion S0 and (S2+S2*) form in Western blots as described in (A).

### SARS-CoV-2 E and M proteins alter maturation of the S glycoprotein

To determine which other viral protein could influence S processing and maturation, we co-expressed S with either E, M or N structural proteins in VeroE6 cells. When we determined the ratio of S0/S2 cleavage, we found a strong reduction of S0 cleavage upon S co-expression with E or M, which almost prevented detection of S2 or S2* (Figure 2A). In contrast, co-expression with N did not influence the processing of S. Of note, we found that the cleavage rate of S was increased in 293T cells as compared to that in VeroE6 cells (Figure 1A, 1B), allowing to better study the influence of E, M or N structural proteins on the regulation of S cleavage and maturation. Hence, we confirmed in 293T cells that co-expression of S with M or E – though not with N – decreased the proportion of S cleaved forms (Figure 2B, 2D). Note that the lower band of S2 (S2*) that was observed in SARS-CoV-2 infected cells (Figure 1A) was detected upon co-expression of S with E or with M (Figure 2B), indicating that both E and M influence maturation of S2 (see below). Finally, we found that the co-expression of E induced a decrease of the total amounts of S detected in lysates of transfected VeroE6 and 293T cells, with a lower impact of E in 293T cells compared to VeroE6 cells (Figure 2A-C), suggesting that E may promote S degradation. This hypothesis is addressed in Figure 4 (see below).

**Figure 2.**
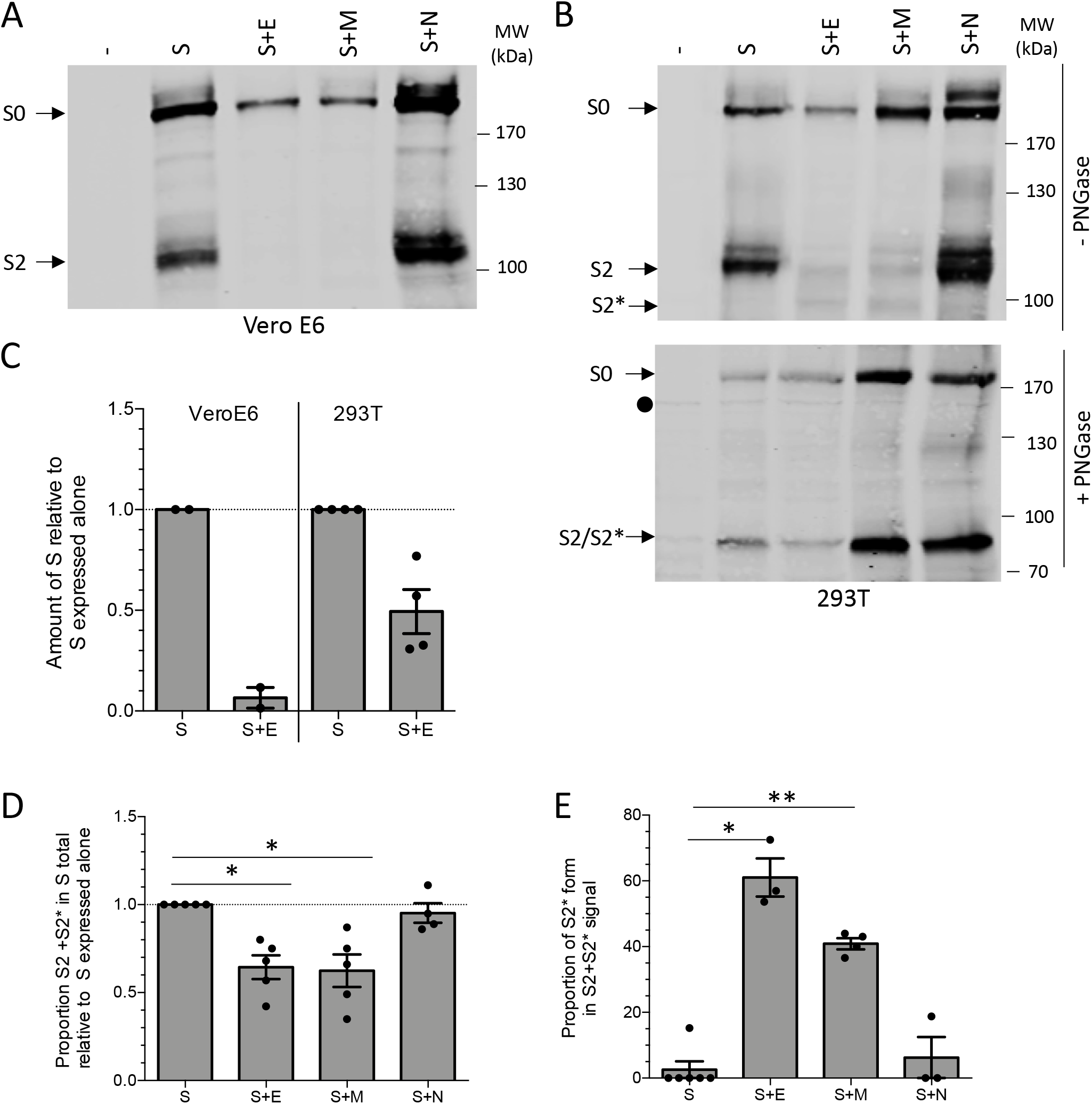
Co-expression of SARS-CoV-2 E and M alter S processing and maturation. **(A)** Representative Western blot of VeroE6 cells transfected with a plasmid encoding S alone or S combined with plasmids expressing E or M or N. The arrows represent S0 and S2 forms. **(B)** Representative Western blot of 293T cells transfected with a plasmid encoding S alone or S combined with plasmids expressing E or M and N. Cells lysates of 293T cells transfected with a plasmid encoding S alone or S combined with plasmids expressing E or M and N, were left untreated (-PNGase) or were treated with PNGase (+PNGase) to remove glycans. The arrows represent S0, S2 and S2* forms. **(C)** Quantification of amount of S (S0+S2+S2*) in 293T cells or VeroE6 cells transfected with a plasmid encoding S alone or S combined with a plasmid expressing E. **(D)** Quantification of (S2+S2*) proportion in western blot analysis as described in (B). **(E)** Quantification of the proportion S2* forms in S2 signal in western blot analysis as described in (B). The black circle represents an unspecific band.

Our above-described results upon S co-expression with E or M highlighted the presence of the second form of S2, S2* (Figures 1A, 2B and quantification in Figure 2E), which migrated with a higher mobility. Since the S protein is highly glycosylated [15], we thought that this could reflect a change in its N-glycan maturation profile. To address this possibility, we treated lysates of S *vs*. S and E, M, or N co-transfected 293T cells with PNGase F, which removes all N-linked oligosaccharides from glycoproteins. After PNGase F treatment, the S2 and S2* bands were resolved in a single band migrating with the same mobility for all the conditions (Figure 2B). This indicated that S2* is a glycosylation variant of S2 and suggested that the presence of E or M alters the N-glycosylation maturation of S.

### Intracellular retention of SARS-CoV-2 S is induced by M and E, and prevents syncytia formation

Since the E and M proteins of other coronaviruses are involved in the regulation of S localization [16] and since furin is predominantly found in the late compartments of the cell secretory pathway [14], we reasoned that the difference of S cleavage rates between infected *vs*. transfected cells could be due to a difference in S intracellular localization. We first investigated the cellular localization of SARS-CoV-2 S expressed alone *vs*. co-expressed with other structural viral proteins compared to full length virus. First, we found that S expressed alone in VeroE6 cells was widely distributed within the cell, including at the cell surface (Figure 3A), whereas S detected in infected VeroE6 cells was mainly retained intracellularly and was found to strongly colocalize with GM130 (Figure 3A, 3B), a marker of the cis-Golgi but also of some compartment closed to ERGIC [17]. Second, through co-transfection of VeroE6 cells, we found that SARS-CoV-2 M or E proteins co-expressed with S induced its retention inside the cells as judged by its increased colocalization with GM130 (Figure 3A, 3B), suggesting that both M or E can alter the localization of S and prevent its expression at the cell surface. Indeed, the above results were confirmed using anti-S antibody staining of transfected cells without permeabilization (Supplemental Figure 1). In contrast, co-expression with the N protein did not impact the subcellular localization of S (Figure 3A, 3B, and Supplemental Figure 1).

**Figure 3.**
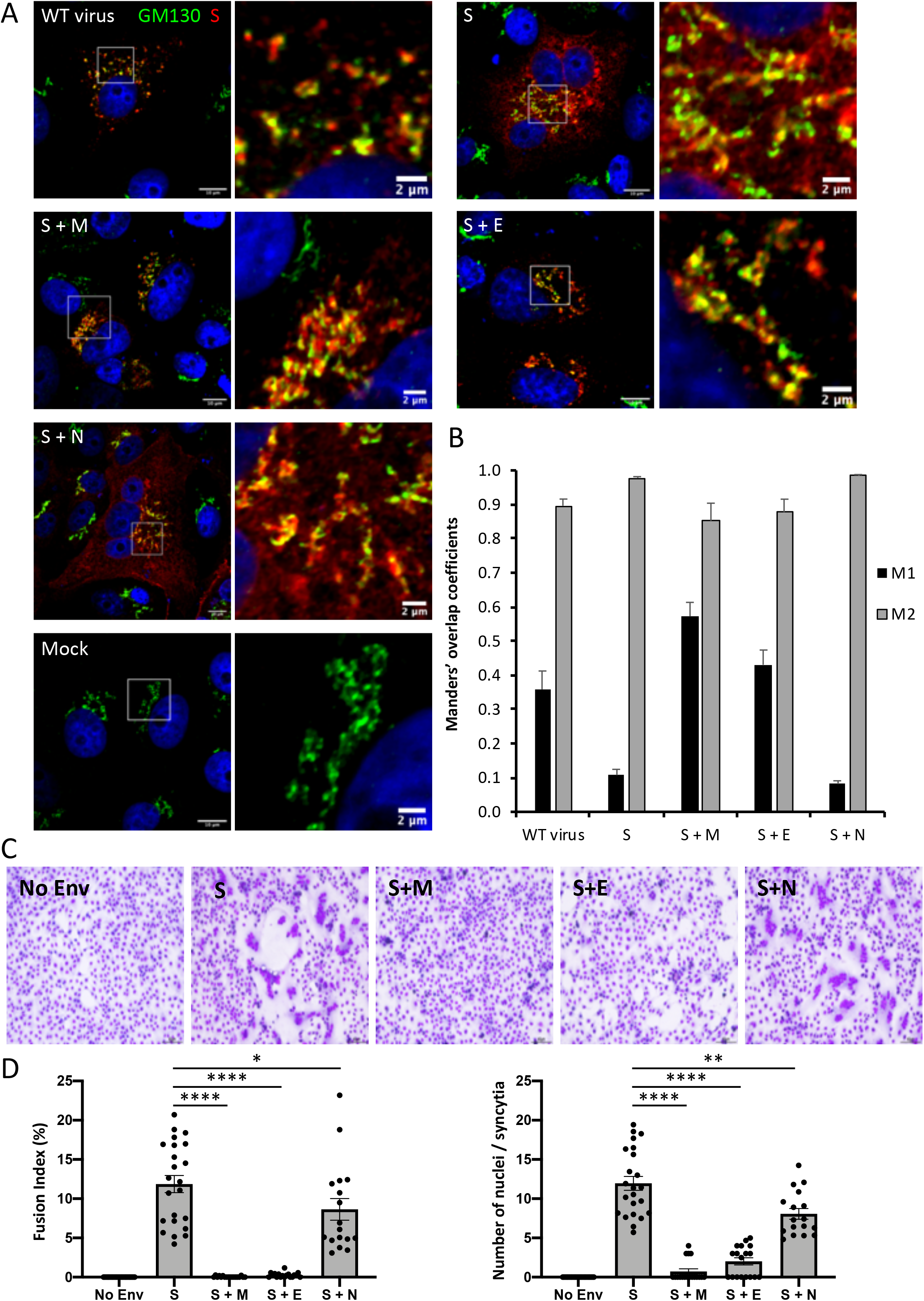
Expression of SARS-CoV-2 E and M induced the retention of S thus preventing cell-cell fusion mediated by S. **(A)** Representative confocal microscopy images of VeroE6 cells infected or transfected with a plasmid encoding S alone or S combined with plasmids expressing M, E or N. The cis-Golgi was revealed with the anti-GM130 antibody (green channel), the S protein was revealed with the anti-SARS-CoV2 S1 antibody (red channel) and the nucleus was revealed with Hoechst. Scale bars of panels and zooms from squared area represent 10μm and 2μm, respectively. **(B)** The Manders’ coefficient M1 represents the fraction of S overlapping with GM130, and the M2 represents the fraction of GM130 overlapping with S. **(C)** Representative pictures of cell-cell fusion assay on VeroE6 cells transfected with a plasmid encoding S alone or S combined with plasmids expressing M, E or N. **(D)** Fusion index (left) and number of nuclei per syncytia (right) of the different conditions as described in (C).

We also found that expression of SARS-CoV-2 S alone induced the formation of syncytia in transfected VeroE6 cells, resulting in the formation in multinucleated giant cells (Figure 3C). This confirmed the presence of S at the cell surface and indicated that all factors required to mediate cell-cell fusion events were present at the surface of these cells. In contrast, we detected strongly reduced and/or much smaller syncytia when S was co-expressed with E or M, whereas S fusion activity was not changed upon N expression (Figure 3C, 3D).

Altogether, these results indicated that M and E regulate the localization of S, probably by allowing its intracellular retention within assembly sites in the ERGIC or cis-Golgi. Since the ERGIC and cis-Golgi are compartments located upstream of organelles in which furin is mainly localized [14], this agrees with a poorer processing and maturation of S upon its co-expression with E and M.

### SARS-CoV-2 E influences the level of expression of S and induces its retention *via* slowing down the secretory pathway

As noted above, our results indicated that E influences the level of expression of S (Figure 2A-C). We thus wondered if E could induce degradation of S *via* the proteasome pathway or, alternatively, *via* lysosomal degradation. To address either possibility, we tested if either MG132, a proteasome inhibitor, or Bafilomcyin A1, an inhibitor of lysosome acidification, could restore the level of S expression. Intriguingly, we found that when expressed alone in 293T cells, S was degraded *via* both the proteasome and the lysosome since treatment of transfected cells with either MG132 or bafilomycin A1 increased S levels by up to 4-6 folds (Figure 4A, 4B). We also found that either drug only slightly increased S levels when co-expressed with E, though to a lesser extent (up to 1.7-fold) than for S individual expression (Figure 4A, 4B), suggesting that E induces the degradation of S in a proteasome- and lysosome-independent manner.

**Figure 4.**
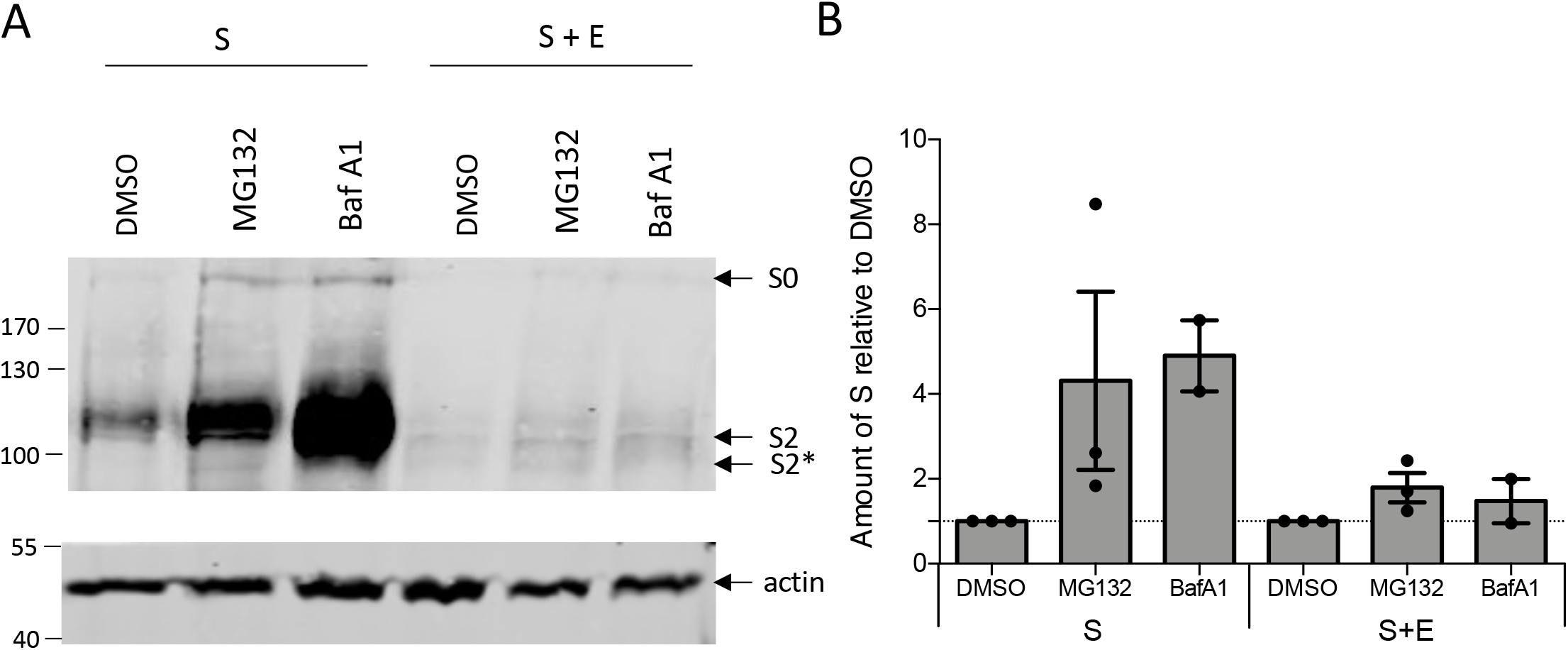
Expression of SARS-CoV-2 E influences S level of expression. **(A)** Representative Western Blot of lysates of 293T cells transfected with a plasmid encoding S alone or S combined with a plasmid expressing E, and treated 16h with MG132 (40uM) or Bafilomycin A1 (100nM) or DMSO as vehicle. The arrows represent S0, S2 and S2* forms and actin used as loading control. **(B)** Quantification of western blot as described in (A).

As shown in Figure 3, co-expression of S with E induced S intracellular retention. As E of some other coronaviruses is supposed to act as a viroporin [4] and as viroporins of alternative unrelated viruses [18–20] have been shown to alter intracellular organelles, we hypothesized that SARS-CoV-2 E could induce the retention of S by slowing down the cell secretory pathway. To demonstrate this, we wondered if E could impact the secretion of VSV-G tsO45 (VSV-Gts), a temperature-dependent folding mutant of VSV-G, a heterologous viral glycoprotein commonly used as model cargo of protein secretion. At 40°C, this protein remains unfolded, resulting in its accumulation in the ER, whereas its folding can be restored at 32°C, which allows its transfer from the ER to the Golgi and then to the plasma membrane.

We transfected Huh-7.5 cells with VSV-Gts in presence of E or of hepatitis C virus (HCV) p7 used as a positive control [18]. First, to address if E alters the traffic from the ER to the *cis*-Golgi, we measured the resistance of intracellular VSV-Gts to endoH digestion [21]. While at 0h, all VSV-Gts glycans remained endoH-sensitive, reflecting ER retention at 40°C, they progressively became resistant to endoH cleavage upon incubation at 32°C for 1h to 3h (Figure 5A, 5B), underscoring VSV-Gts transfer to the Golgi apparatus. We noticed that E expression induced a dose-dependent decrease of the kinetics of VSV-Gts endoH-resistance acquisition (Figure 5B). We confirmed these results in transfected VeroE6 cells (Figure 5C). Interestingly, these latter cells have a lower trafficking speed compared to Huh 7.5 (compare control conditions in Figure 5C); yet, in both cases, E was able to slow down their cell secretory pathway. Next, to address the influence of E on trafficking to the plasma membrane, we analyzed the accumulation of VSV-Gts at the cell surface after incubation of transfected cells at 32°C for different times (Figure 5D). As monitored by flow cytometry analysis, E expression significantly reduced the kinetics and levels of VSV-Gts cell surface expression (Figure 5D, 5E).

**Figure 5.**
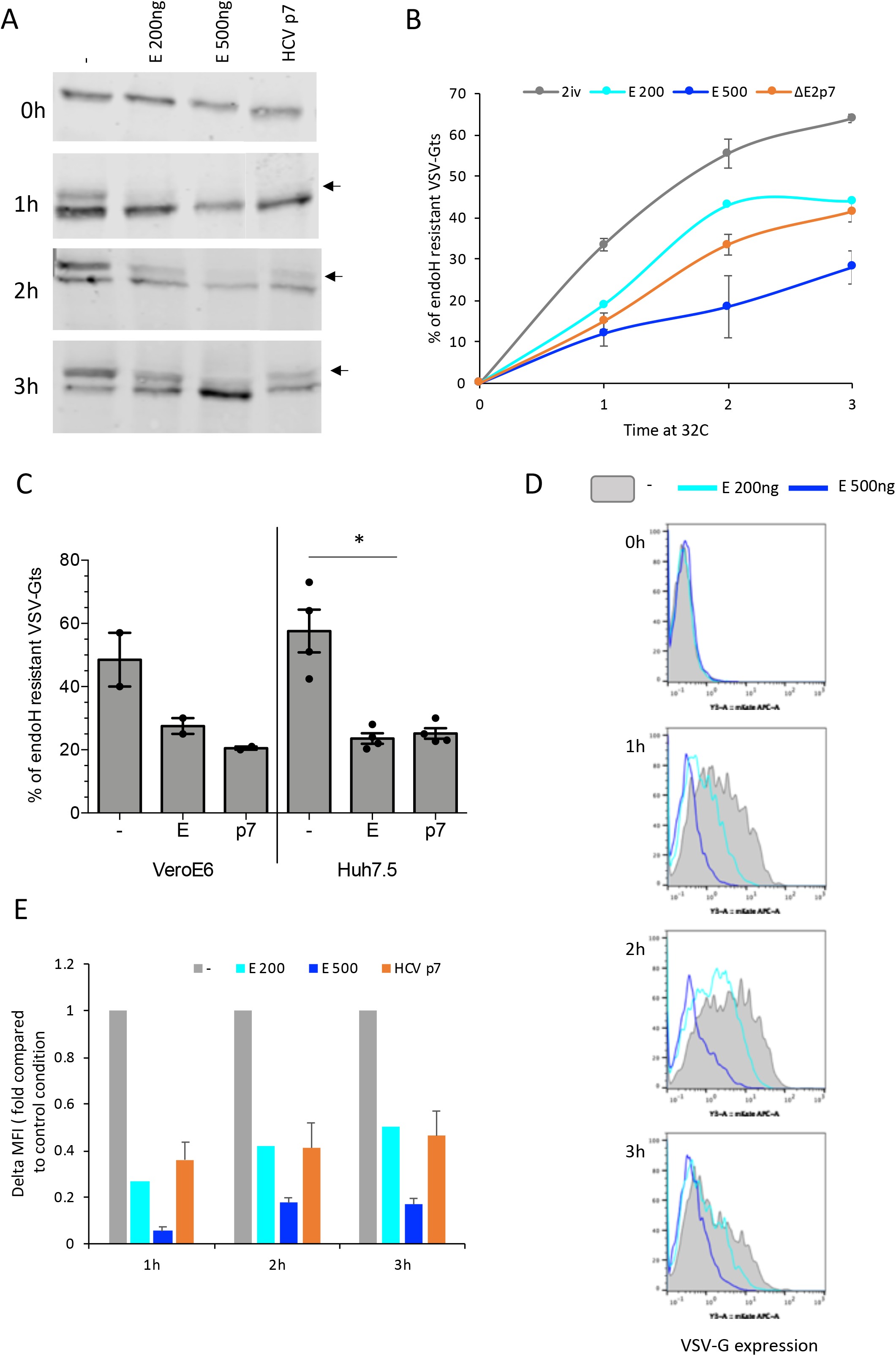
SARS-CoV-2 E induces retention of S *via* slowing down the cell secretory pathway. Huh7.5 or VeroE6 cells were transfected with vectors encoding VSV-Gts and HCV p7 (JFH1) or SARS-CoV E at two different doses, as indicated. Transfected cells were grown overnight at 40°C, which maintains VSV-Gts unfolded and results in its accumulation in the ER. Cells were then incubated for different periods of time (0h, 1h, 2h and/or 3h, as indicated) at 32°C, which allows restoration of its folding and thus, its secretion. **(A)** Representative western blot analysis of cell lysates co-expressing VSV-Gts and E or p7, digested with endoH glycosidase. The endoH-resistant VSV-Gts species (arrows) indicate proteins that traffic to and beyond the Golgi apparatus **(B)** Quantification of western blots as described in (A). **(C)** Quantification of western blot analysis of cell lysates of Huh7.5 or VeroE6 cells co-expressing VSV-Gts and E or p7, lysed at 3h (VeroE6 cells) or 2h (Huh7.5) post-temperature shifting and digested with endoH glycosidase. The timing was chosen to have the same percentage of endoH resistant forms of VSV-Gts in both cell types. **(D)** Representative histogram of cell surface expression of VSV-Gts assessed by flow cytometry, using the 41A1 mAb directed against VSV-G ectodomain. Gray plot represented control cells, light blue line cells with E at 200ng, and dark blue line cells with E at 500ng. **(E)** Cell surface expression of VSV-Gts assessed by the variations of the mean fluorescence intensity (delta MFI) of cell surface-expressed VSV-Gts relative to time 0h at 32°C.

Altogether, these results indicated that SARS-CoV-2 E protein slows down the cell secretory pathway, hence inducing a non-specific retention of glycoproteins, which also includes S.

### The C-terminal moiety of SARS-CoV-2 S cytoplasmic tail is essential for M-mediated retention of S

Previous studies showed that for SARS-CoV-1 S protein, a dibasic retrieval signal KxHxx present at the C-terminus of its cytoplasmic tail allows the recycling of S *via* binding to COPI [12]. Such a recycling of S increases its capacity to interact with M, which resides at the virion assembly site. Owing to the conservation of this motif in the cytoplasmic of SARS-CoV-2 S (Figure 6A), we thought to investigate if the involved mechanism is conserved. Therefore, we tested the impact of M on retention of a mutant of SARS-CoV-2 S, named SΔ19, from which the last 19 amino acids, including the dibasic retrieval signal, were removed (Figure 6A). In contrast to wt S, we found that SΔ19, when co-expressed with M in VeroE6 cells, exhibited impaired intracellular retention (compare Figure 6B with Figure 3B, and Supplemental Figure 1), which confirmed that this retrieval signal allows S recycling and consequently, M-mediated retention of S. Interestingly, co-expression of E still induced the retention of SΔ19 (Figure 6B), which agreed with our above results that E can induce the retention of S by modulating intracellular trafficking (Figures 3 and 5) rather than by interacting directly with S. Of note, we observed that despite the presence of E, SΔ19 did not colocalize with GM130 to the same extent than for S (compare M1 coefficients in Figure 6B vs. Figure 3B), suggesting that E induces the retention of SΔ19 inside the cells though, due to the lack of retrieval signal, SΔ19 does not accumulate in GM130-containing compartments.

**Figure 6.**
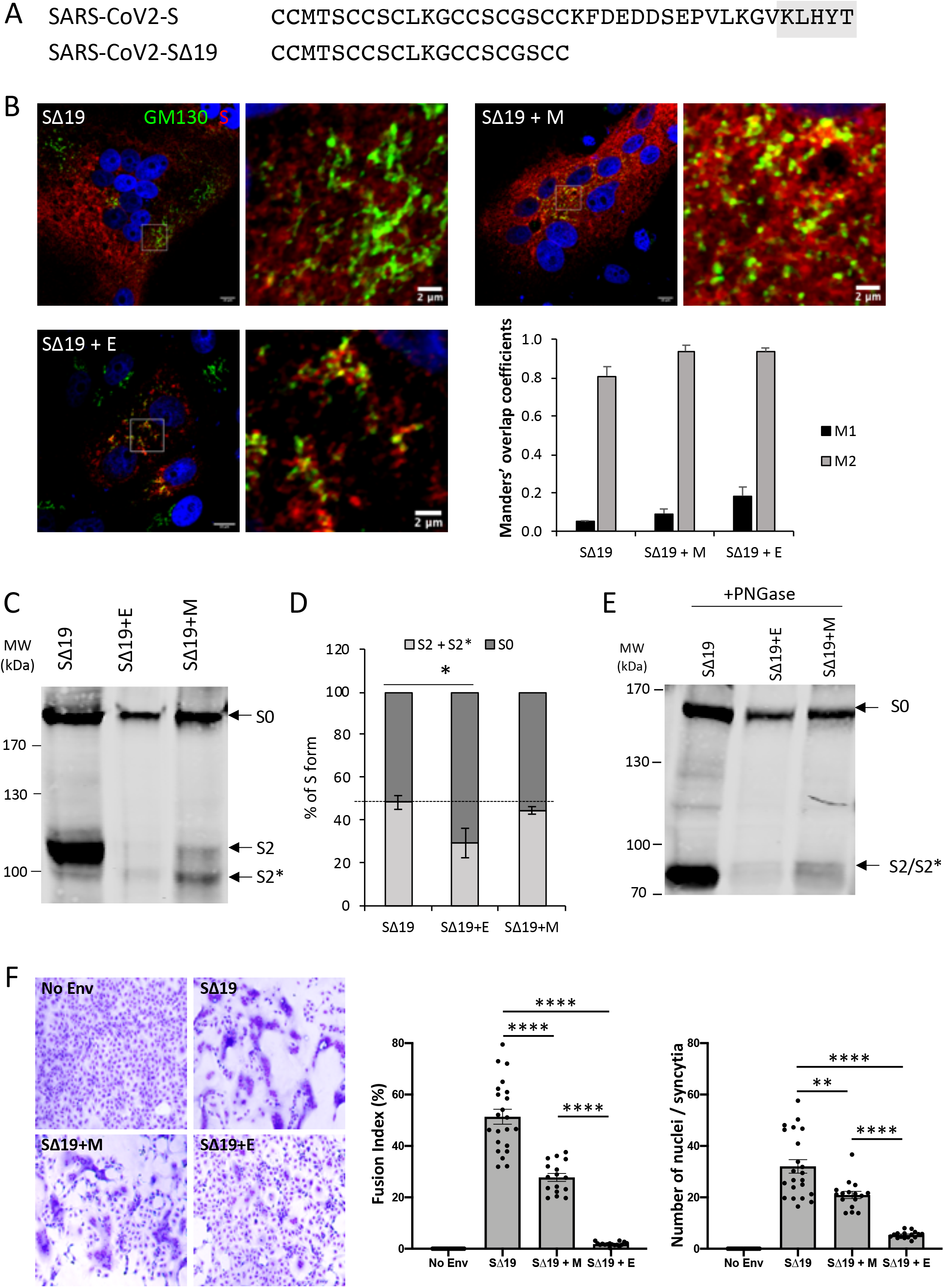
The C-terminal moiety of S cytoplasmic tail is essential for M-mediated retention of SARS-CoV-2 S. **(A)** Alignment of sequences of the last amino-acids of S of SARS-CoV-2 or mutated by deletion of the last 19 amino acids (SΔ19). **(B)** Representative confocal microscopy images of VeroE6 cells transfected with a plasmid encoding SΔ19 alone or SΔ19 combined with plasmids expressing M or E. The cis-Golgi was revealed with the anti-GM130 antibody (green channel), the S protein was revealed with the anti-SARS-CoV2 S1 antibody (red channel) and the nucleus was revealed with Hoechst. The Manders’ coefficient M1 represents the fraction of S overlapping with GM130, and the M2 represents the fraction of GM130 overlapping with S. Scale bars of panels and zooms from squared area represent 10μm and 2μm, respectively. **(C)** Representative Western Blot of 293T transfected with a plasmid encoding SΔ19 or SΔ19 combined with plasmids encoding E or M. The arrows represent S0, S2 and S2* forms. **(D)** Quantification of S form of independent western blot as described in (C). **(E)** Cells lysates of 293T cells transfected with a plasmid encoding SΔ19 alone or SΔ19 combined with plasmids expressing E or M and N were treated with PNGase F to remove glycans. The arrows represent S0, S2 and S2* forms. **(F)** Representative pictures of cell-cell fusion assays on VeroE6 cells transfected with a plasmid encoding SΔ19 alone or SΔ19 combined with plasmids expressing M or E (left). Fusion index and number of nuclei per syncytia of the different conditions (right).

To confirm the correlation between the lack of retention and processing of S, we determined the cleavage rate of SΔ19 in cells co-transfected with E or M. SΔ19 co-expressed with E exhibited reduced cleavage rate whereas co-expression of M did not alter this rate (Figure 6C, 6D). This confirmed that the M-mediated retention of S and its reduced cleavage rate is dependent on the C-terminal retention motif whereas E-mediated retention of S and its reduced cleavage rate is linked to modification of the cell secretory pathway.

To corroborate these results, we determined the fusion index of cells expressing SΔ19 alone or SΔ19 in combination with the other viral structural proteins. Interestingly, we found that SΔ19 was highly fusogenic and induced much larger syncytia than wt S (Figure 3D *vs*. Figure 6F), likely because of its accumulation at the cell surface owing to deletion of the recycling signal. In agreement with our previous observations that M, but not E, does not alter SΔ19 processing (Figure 6C, 6D), co-expression of SΔ19 and M did not change the fusion index whereas co-expression of SΔ19 and E almost suppressed the formation of syncytia (Figure 6F).

Interestingly, we detected the presence of both S2 and S2* for SΔ19 co-expressed with E or M (Figure 6C), which was resolved to a single S2 band upon treatment with PNGase F (Figure 6E), thus confirming that, for both wt S and SΔ19, the presence of S2* was due to modification of N-glycan maturation. Since M is not able to induce retention of SΔ19, this argues for a modification of N glycosylation pathway by E and M independently of the retention of S.

Altogether, these results indicated that M and E induce the retention of S *via* different mechanisms. Indeed, the cytoplasmic tail of S may be involved in its weak retrieval by allowing interaction with COPI, and thus a subsequent interaction with M, whereas E may induce the retention of S by regulating intracellular trafficking.

### Secretion of S-displaying VLPs requires co-expression of both E, M and N

Previous reports indicated that for alternative coronaviruses, the intracellular retention of S by M is essential for assembly of infectious particles and that the presence of E is essential for budding of particles. We sought to extend this notion by investigating the mechanism of assembly of SARS-CoV-2 VLPs, which appears poorly defined. Since we showed that M and E are involved in the regulation of S localization and trafficking (Figure 3), we hypothesized that either protein could be required for production of VLPs. Thus, we transfected 293T cells with plasmids inducing expression of S alone *vs*. of S in combination with E, M, and/or N or with all proteins altogether. At 48h post-transfection, we collected the cell supernatants and we purified particles by ultracentrifugation through a sucrose cushion. As shown in Figure 7A-C, we found that S expressed alone, *i.e*., raising S0 and S2 bands, was poorly detected in the pellets of ultracentrifugation. Co-expression of N, M or E alone with S did not improve the secretion of S. Co-expression of S with both E and N or with both M and N could slightly increase the presence of S in the pellets (Figure 7A, 7B), though this correlated to an increased expression level in cell lysates (Figure 7C). In contrast, we found that co-expression of the combination of M, N and E with S induced a strong production of VLPs with a high detection of S in the pellet and low detection in the cell lysate, suggesting that all structural proteins are required for optimal secretion of S-containing VLPs (Figure 7A-C). We also found that N expressed with S was poorly secreted, whereas its secretion was readily increased upon co-expression with S, M and E (Figure 7A), hence suggesting a concerted action of N, M and E for budding and secretion of S-containing VLPs.

**Figure 7.**
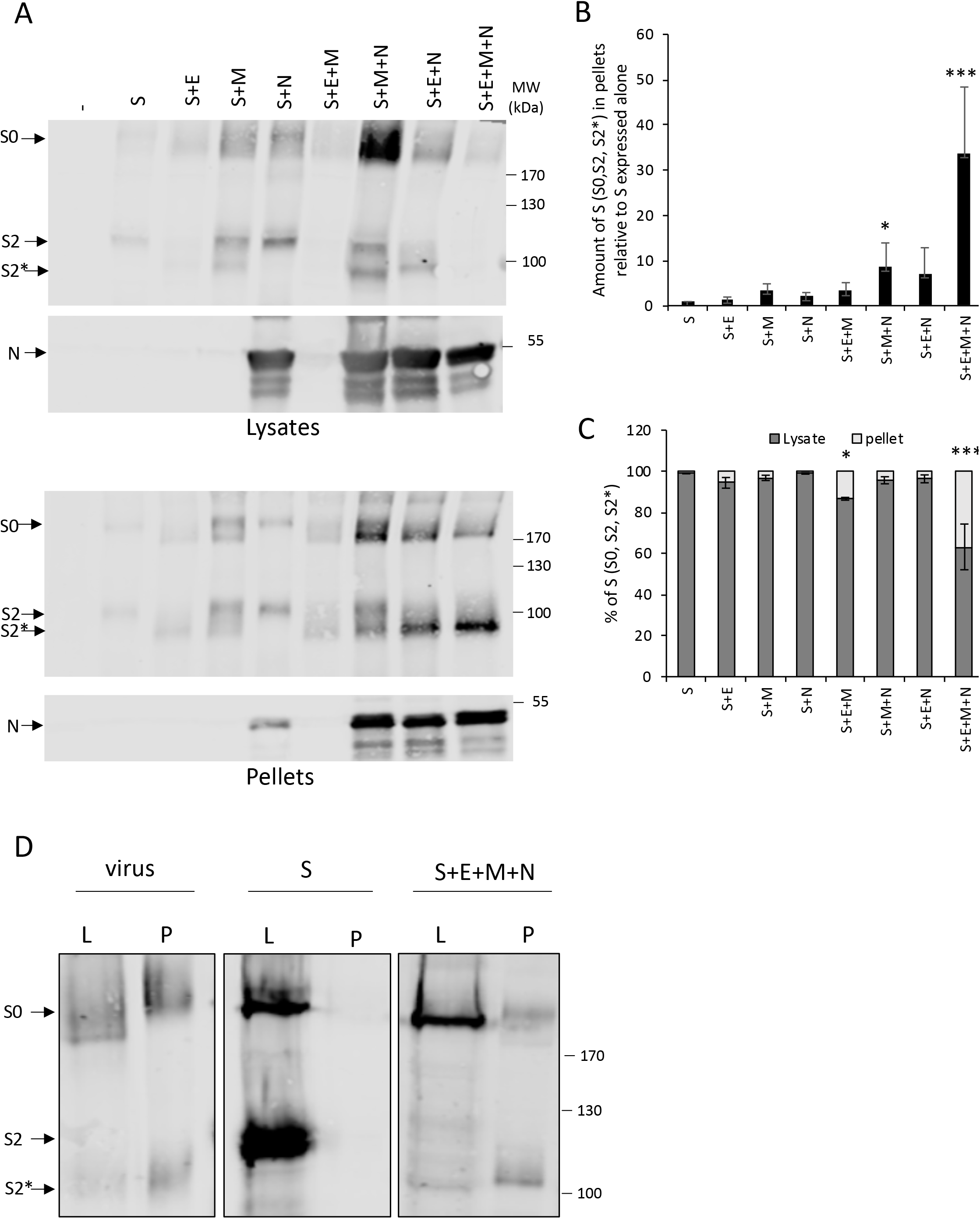
Secretion of SARS-CoV-2 S-displaying VLPs requires expression of both E, M and N. **(A)** Representative western blot of cells lysates and pellet of 293T transfected with a plasmid encoding S alone or S combined with plasmids encoding E, M and N. The arrows represent S0, S2 and S2* forms and N. **(B)** The amounts of S detected in pellets of ultracentrifugated supernatants of producer cells were determined by quantification of independent western blots as described in (A). **(C)** Proportion of S in lysates and pellets determined by quantification of independent western blot as described in (A). **(D)** Western Blot analysis of cell lysates (L) or pellets (P) or VeroE6 cells infected with full-length virus or transfected with S alone or with S, E, M and N expressing plasmids. The arrows represent S0, S2 and S2* forms.

Similar to our observations in lysates of co-transfected cells (Figure 1), we found that the S2* form was detected in the pellets of purified particles produced upon S co-expression with E or M, as compared to those produced with S alone or with S and N (Figure 7A), in agreement with a different maturation pathway when E or M are present, as above-proposed.

Finally, we confirmed that expression of S, E, M and N in VeroE6 cells allowed the secretion of VLPs (Figure 7D). We observed the presence of S in the pellets of supernatants of either infected cells or cells transfected with S, E, M and N. Note that we confirmed the presence of the S2* form in the latter pellets.

Altogether, these results showed that E, M, and N are required for the optimal production of VLPs containing S.

## Discussion

Here, we highlight that SARS-CoV-2 E and M proteins induce the retention of S inside the cells, which probably provides a mechanism allowing the targeting of S close to the virion assembly site. In addition, we also show that independently of their effect on retention of S, E and M co-expression alter the maturation of the N glycans of S (Figure 8). Finally, we found that E, M and N are the optimal set of proteins required to for secretion S-harboring virus-like particles.

**Figure 8.**
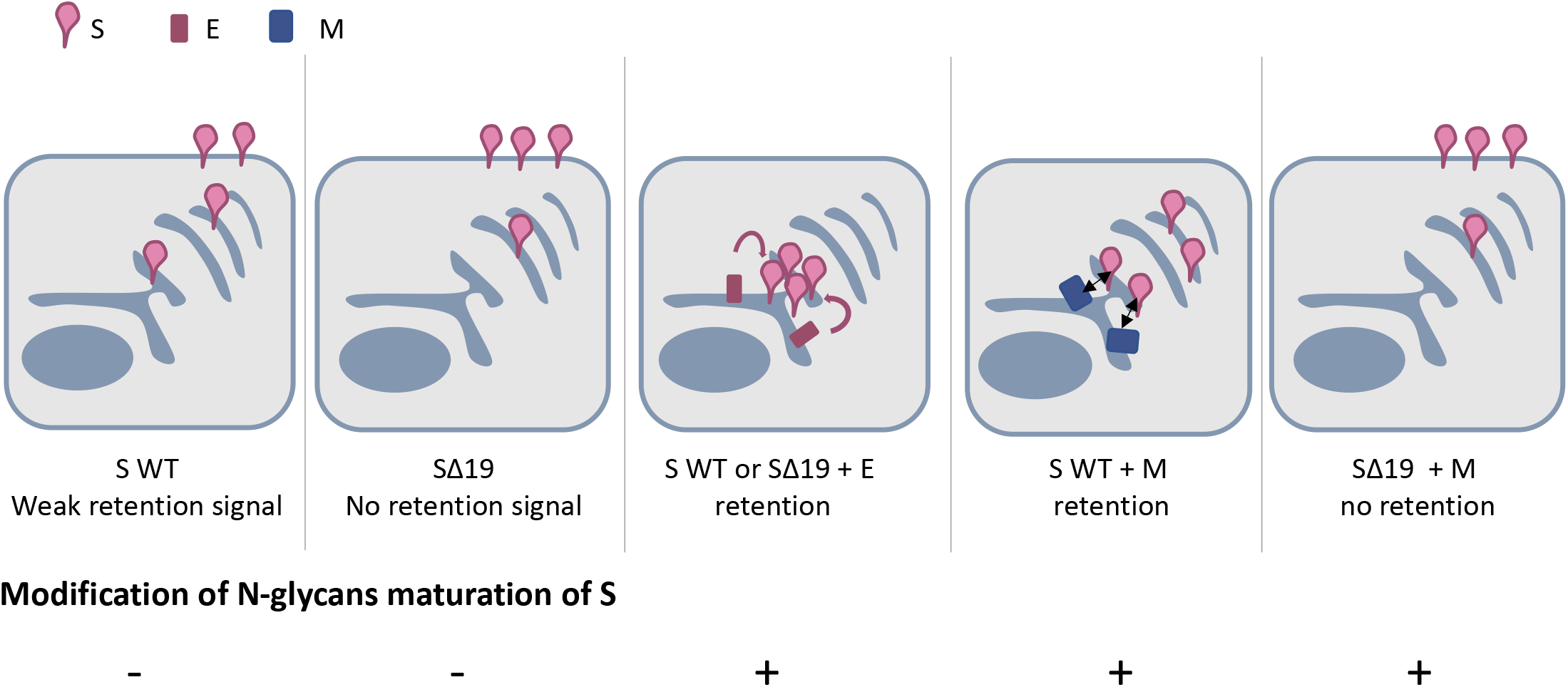
Model of localization of SARS-CoV2 S protein. Due to its weak retention signal located at the C-terminus of its cytoplasmic tail, S is found at the cell surface but also inside the cells. In contrast, removal of the last 19 amino acids (SΔ19) increases the presence of S at the cell surface. Co-expression of E induces the retention of both wt S and SΔ19. In contrast, co-expression of M induces the retention of wt S only. Irrespective of S forms, the presence of E and M modulate the maturation of N-glycans of S.

### SARS-CoV-2 E slows down the cell secretory pathway

We showed that E induce the retention of S by modulating the cell secretory pathway (Figure 5). Interestingly, E from other β-coronaviruses, and especially SARS-CoV-1, were shown to form cation-selective ion channels [22, 23]. We and others previously demonstrated that viroporins of unrelated viruses are able to slow down the cell secretory pathway, like for HCV p7 [18] or Influenza A virus M2 [19, 20], prompting us to investigate if SARS-CoV-2 E could alter the cell secretory pathway. In addition, as it was shown that Coronavirus Infectious Bronchitis Virus (IBV) E is able to alter the secretory pathway [24], we therefore speculate that SARS-CoV2 E could alter the cell secretory pathway *via* a mechanism shared with some other coronaviruses. Interestingly, we also found that the retention of S induced by E is independent of the retrieval signal of S (Figure 6), although this retrieval signal modulates S localization even in presence of E. Indeed, in presence of E, a mutant of S deleted for this signal did not accumulate in GM130-containing compartment in contrast to wt S.

We propose that the modulation of the cell secretory pathway could be important for assembly of infectious particles by allowing the accumulation of the viral structural components at the virion assembly site. Alternatively, the modulation of cell secretory pathway *per se* could be independent of the assembly of infectious viral particles, but rather linked to virulence and/or induction of inflammasome since E was found to be associated to virulence of several coronavirus genera, like *e.g*., for SARS-CoV-1 [25, 26] or IBV [24] as well as induction of inflammasome for SARS-CoV-1[27].

### SARS-CoV-2 E influence the level of expression of S

We found that expression of E decreased the level of expression of S (Figure 2). One possibility is that E could induce a degradation of S *via* either the proteasome or the lysosome pathways. However, inhibition of the latter pathways did not restore the level of S expression in presence of E, hence suggesting that E does not act on the degradation of S *per se* but rather, could alter S synthesis and/or modify the level of secretion of S. Yet, we ruled out the latter hypothesis since we did not observe an increase of secretion of S in presence of E (Figure 7), hence, inferring that E may regulate the synthesis of S. Indeed, E may induce cellular stress *via* its ion channel activity, as proposed for some other viroporins [28], which may thus influence the level of translation by cellular stress responses [29]. In line with this hypothesis, individual expression of E from SARS-CoV-1 [30] as well from the murine coronovirus MHV (mouse hepatitis virus) [31] could induce apoptosis. However, we cannot rule out that the effect observed here for E of SARS-CoV2 could be due to the overexpression of E alone, as compared to the expression E in the context of infection, since the expression of other viral proteins could compensate the effect of E.

### Expression of SARS-CoV-2 E and M modulate the N-glycosylation pathway

We found that E and M regulate the maturation of N glycosylation of S (Figure 2); yet, we showed that this is not related to the role of the former proteins in the retention of S at the Golgi, as assessed by using the SΔ19 mutant that retained the same maturation as wt S despite its lack of intracellular retention (Figure 6). Rather, this suggested that the modification of N-glycosylation is not linked to glycoprotein retrieval in an intracellular compartment lacking the glycosyltransferases. Previous reports have shown that, for other coronaviruses, E and M are located at the ERGIC and/or Golgi membranes [32–35]. Although we could not confirm this for SARS-CoV-2 E and M, owing to the unavailability of specific antibodies, we speculate that they share the same intracellular localization. Since the maturation of N-glycosylation occurs in the Golgi, one possibility is that accumulation of E and M proteins at the membrane of this organelle could induce changes that alter the correct action of glycosyltransferases [36].

### All structural proteins are required for optimal production of SARS-CoV-2 VLPs

While S expressed alone did not induce the secretion of S-containing VLPs, we found that combining its expression with some of the other structural proteins resulted in the formation of such particles, although co-expression of all structural proteins was the most efficient combination to induce VLP secretion (Figure 7). Indeed, when all these proteins were expressed, the cells were almost depleted in S whereas S was readily found in the pellets of their ultracentrifugated supernatants (Figure 7A). While M is essential for the assembly of virions, previous results of others showed that for alternative coronaviruses, S is dispensable for promoting virion assembly although it is readily incorporated in viral particles upon co-expression with other structural proteins [16]. Thus, we could speculate that SARS-CoV-2 has adopted a similar mechanism. However, the involvement of E and N in virion assembly might be coronavirus strain-dependent [16]. For SARS-CoV-1, the mechanism of formation of VLPs remains unclear since co-expression of M and E [37], of M and N [8], or of M, N and E [38] proteins resulted in the production of VLPs, although these previous studies did not focus on S-containing VLPs. Our results underscore that for SARS-CoV-2, S co-expression with either E, M, or N structural proteins induces low level secretion of S containing-VLPs. We also tested all combination of the three structural proteins E, M and N co-expressed with S. While all conditions allowed secretion of S containing-VLPs, we found that the optimal VLP production required both M, N and E co-expression.

Previous results indicated that for most coronaviruses, E is essential for incorporation of M in viral particles, inferring a conserved mechanism for SARS-CoV-2. First, E and M are known to interact with each other [33]. In addition, E might be involved in inducing membrane curvature or scission of vesicles [39, 40]. The role of N is more complex and remains poorly defined. N is able to form high-order oligomers [41, 42] even in absence of RNA [43] that may stabilize the particles and or the oligomerization of M. In addition, we showed that N can be secreted in presence of S independently of E and M expression, suggesting that N, at least in presence of S, may help virion budding *via* a “push” mechanism [44]. Indeed, the driving force for budding of enveloped viruses can be driven by the nucleocapsid itself that “pushes” a membranous bud, *via* specific inner structural proteins (*e.g*., Gag precursor of HIV), or alternatively, by the envelope glycoproteins that can form a symmetric lattice “pulling” the membrane (*e.g*., prME of flaviviruses), even if viruses have evolved and developed different mechanisms with some variations or combinations between these two main models [44]. In line with this, we could imagine that for SARS-CoV-2, the optimal driving force for budding could be due to N that could push the membrane as well as to E and M that could create optimal curvature and pull the membrane, hence allowing efficient budding of viral particles.

Altogether, the results of this report indicated that E and M proteins differentially influence the properties of S proteins to promote assembly of SARS-CoV-2VLPs. Our results therefore highlight both similarities and dissimilarities in these events, as compared to other β-coronaviruses. Owing to their lack of infectivity, such VLPs could provide attractive tools for studying vaccines or immune responses against COVID-19.

## Materials and Methods

### Cell culture and reagents

Huh7.5 cells (kind gift of C. Rice, Rockefeller University, New York, USA), Vero E6 cells (ATCC CRL-1586) and 293T kidney (ATCC CRL-1573) cells were grown in Dulbecco’s modified minimal essential medium (DMEM, Invitrogen, France) supplemented with 100U/ml of penicillin, 100μg/ml of streptomycin, and 10% fetal bovine serum.

### Plasmids

Homo sapiens codon optimized SARS-CoV-2 S (Wuhan-Hu-1, GenBank: QHD43419.1) was cloned into pVAX1 vector. The delta 19 truncation of S form was generated by site directed mutagenesis introducing a stop codon after Cys1254 [45]. SARS-CoV-2 E, M and N genes (Wuhan-Hu-1, GenBank: QHD43419.1) were synthesized and cloned into pCDNA3.1(+) vector. The plasmid pEGFP-N3-VSV-Gts (kind gift from K. Konan, Albany Medical College, USA). The plasmids encoding HCV ΔE2p7(JFH1) was described previously [18].

### Antibodies

Mouse anti-actin (clone AC74, Sigma-Aldrich), rabbit anti-SARS-CoV2 S2, mouse anti-SARS-CoV2 S1 and mouse anti-SARS-CoV2 N (Sino Biological), mouse anti-GFP (Roche), anti-VSV-G 41A1 and rabbit anti-GM130 (clone EP892Y, Abcam) were used according to the providers’ instructions.

### Viral production and infection

SARS-CoV-2 particles (kind gift of B. Lina, CIRI, Lyon) are referenced in GISAID EpiCoVTM database (reference BetaCoV/France/IDF0571/2020, accession ID EPI_ISL_411218) and were amplified on VeroE6 cells [46]. Briefly, for stock production, cells were infected with MOI=0.01 in DMEM for 90min at 37°C. Then, medium was replaced with DMEM-2%FCS. Supernatant fluids were collected after two days at 37°C, clarified by centrifugation (400xg, 5min), aliquoted and titrated in plaque forming unit by classic dilution limit assay on the same Vero E6 cells. Lysis and pellet were done as described below.

### VSV-Gts analysis

Huh7.5 cells were seeded 16h prior to transfection with pEGFP-N3-VSV-Gts and p7-encoding plasmid using GeneJammer transfection reagent (Agilent). Medium was changed 4h post-transfection and cells were incubated overnight at 40°C. 24h post-transfection, cells were chased at 32°C. For western blot analysis, cells were lysed at indicated time points in wells cooled on ice before clarification, Endoglycosidase Hf treatment and western blot analysis. Endo-Hf (NEB) treatment was performed according to the manufacturer’s recommendations. Briefly, protein samples were mixed to denaturing glycoprotein buffer and heated at 100°C for 5 min. Subsequently, 1,000 units of Endo-Hf were added to samples in a final volume of 25 μL and the reaction mixtures were incubated for 1 h at 37°C. For flow cytometry analysis, cells were harvested and put in suspension at 32°C. At indicated time points, cells were fixed with 3% paraformaldehyde.

### Analysis of expression different proteins in cell lysate and pellet

HEK293T cells were seeded 24h prior to transfection with the different plasmids (2μg of each plasmid for a 10cm dish) using calcium phosphate precipitation. VeroE6 cells were seeded 24h prior to transfection with the different plasmids (2μg of S, 0.2μg of E, 0.4μg of M and 0.8μg of N) using GeneJammer transfection reagent (Agilent). Medium was replaced 16h post-transfection. Supernatants and cell lysate were done 24h later. Cell were counted and 100,000 cells were lysed in 100μL lysis buffer (20 mM Tris [pH 7.5], 1% Triton X-100, 0.05% sodium dodecyl sulfate, 150nM NaCl, 5% Na deoxycholate) supplemented with protease/phosphatase inhibitor cocktail (Roche) and clarified from the nuclei by centrifugation at 13,000×*g* for 10 min at 4°C for quantitative western blot analysis (see below). For purification of particles, supernatants were harvested and filtered through a 0.45μm filter and centrifuged at 27,000 rpm for 3h at 4°C with a SW41 rotor and Optima L-90 centrifuge (Beckman). Pellets were resuspended in PBS prior to use for western blot analysis.

### Deglycosylation with PNGase F

PNGase F (NEB) treatment was performed according to the manufacturer’s recommendations. Briefly, protein samples were mixed to denaturing glycoprotein buffer and heated at 100°C for 5 min. Subsequently, 20 units of PNGase F were added to samples in a final volume of 25μl with NP-40 and buffer and the reaction mixtures were incubated for 1 h at 37°C, before western blot analysis.

### Western blot analysis

Proteins obtained in total lysates or after digestion were denatured in Laemmli buffer at 95°C for 5min and were separated by sodium dodecyl sulfate polyacrylamide gel electrophoresis, then transferred to nitrocellulose membrane and revealed with specific primary antibodies, followed by the addition of Irdye secondary antibodies (Li-Cor Biosciences). Signals were quantitatively acquired with an Odyssey infrared imaging system CLx (Li-Cor Biosciences).

### Immuno-fluorescence (IF) and confocal microscopy imaging

Immuno-fluorescence experiments were done as previously described [47]. Briefly, 3×10e5 VeroE6 cells grown on coverslips were infected with wt virus (MOI=0.01) or transfected with 1μg of each expressing construct with GeneJammer according to the manufacturer’s instructions. Six hours later, the media of transfected cells was replaced by fresh media and cells were cultured for an additional 18 hours. Twenty for hours post-infection or - transfection, cells were fixed for 15min with 3% PFA and permeabilized or not with 0.1% Triton X-100. After a saturation step with 3% BSA/PBS, cells were incubated for 1 hour with rabbit anti-GM130 and mouse anti-SARS-CoV2 S1 antibodies at 1/200 dilution in 1% BSA/PBS, washed 3 times with 1%BSA/PBS, and stained for 1 hour with donkey anti-rabbit AlexaFluor-488 and donkey anti-mouse AlexaFluor-555 secondary antibodies (Molecular Probes) diluted 1/2,000 in 1% BSA/PBS. Cells were then washed 3 times with PBS, stained for nuclei with Hoechst (Molecular Probes) for 5 min, washed and mounted in Mowiol (Fluka) before image acquisition with LSM-710 or LSM-800 confocal microscopes.

Images were analyzed with the ImageJ software (imagj.nih.gov) and the Manders’ overlap coefficients were calculated by using the JACoP plugin.

### Cell-cell fusion assay

The cell-cell fusion assay was adapted from [48]. Briefly, 3×10e5 VeroE6 cells were transfected with 1μg of the different expression constructs with GeneJammer according to the manufacturer’s instructions. After 6 hours, the transfection media was removed and replaced by fresh media for an additional 24 hours. Thirty hours post-transfection, transfected cells were fixed and counterstained with May-Grünwald and Giemsa solutions (Sigma-Aldrich) according to the manufacturer’s instructions. Between 17 and 24 fields were acquired in 3 independent experiments and the fusion index of the different combinations was determined as (N – S)/T x 100, where N is the number of nuclei in the syncytia, S is the number of syncytia, and T is the total number of nuclei counted.

### Statistical analysis

Significance values were calculated by applying the Kruskal-Wallis test and Dunn’s multiple comparison test using the GraphPad Prism 6 software (GraphPad Software, USA). For fusion index, a two-tailed, unpaired Mann-Whitney test was applied. P values under 0.05 were considered statistically significant and the following denotations were used: ****, P≤0.0001; ***, P≤0.001; **, P≤0.01; *, P≤0.05; ns (not significant), P>0.05.

## Acknowledgments

We thank to Konan Kouacou for the VSV-Gts construct and Charles Rice for the Huh7.5 cells. We also thank B. Lina for the SARS-CoV-2 particles. We thank Lu LU and Moumita MONDAL for technical help in the generation of SARS-CoV2 derived plasmids

We acknowledge the contribution of SFR Biosciences (UMS3444/CNRS, US8/Inserm, ENS de Lyon, UCBL) facilities: LBI-PLATIM-Microscopy, ANIRA-Cytometry for excellent technical assistance and support. We than Didier Décimo for support with the BSL3 facility.

## Fundings

FLC received financial support from the LabEx Ecofect (ANR-11-LABX-0048) of the “Université de Lyon”, within the program “Investissements d’Avenir” (ANR-11-IDEX0007) operated by the French National Research Agency (ANR), the ANR (grant from RA-Covid-19), and the Fondation pour la Recherche Médicale (FRM).

DL received financial support from National Key R&D Program 376 of China (2020YFC0845900, D.La), Shanghai Municipal Science and Technology Major Project (20431900402, D.La).the National Natural Science Foundation of China (31870153 D.La), Chinese academy of Sciences PIFI program (D.La).

The funders had no role in study design, data collection and analysis, decision to publish, or preparation of the manuscript.

---

**Supplemental Figure 1.**
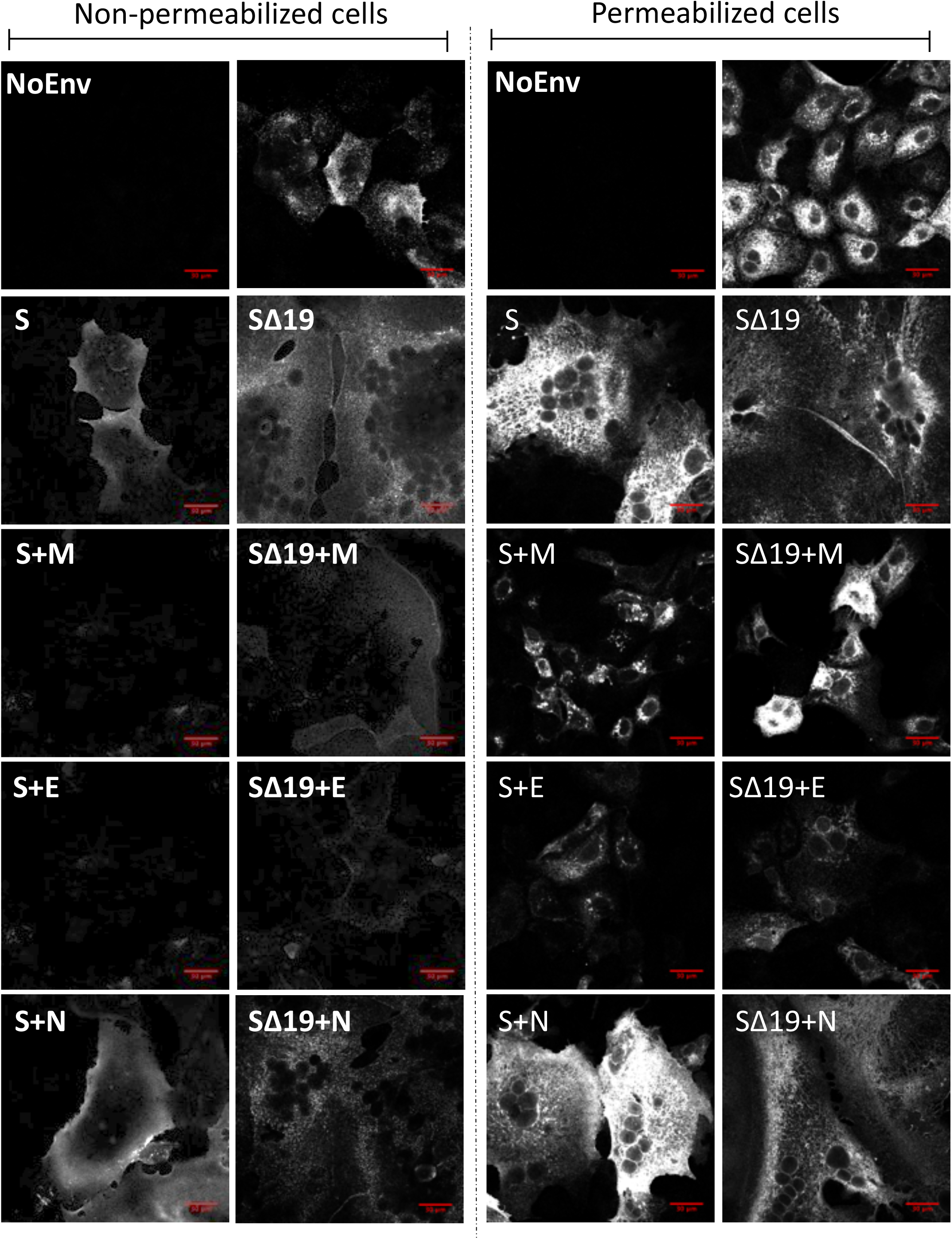
Expression of SARS-CoV-2 E and M reduce the presence of S at the cell surface. Representative confocal microscopy images of VeroE6 cells transfected with plasmids encoding S *vs*. SΔ19 alone or combined with plasmids expressing M, E or N. Cells were fixed and permeabilized, or not, with Triton X-100.

## Notes

### Competing Interest Statement

The authors have declared no competing interest.

